# Deep learning predicts non-coding RNA functions from only raw sequence data

**DOI:** 10.1101/2020.05.27.118778

**Authors:** Teresa M.R. Noviello, Michele Ceccarelli, Luigi Cerulo

## Abstract

Non-coding RNAs (ncRNAs) are small non-coding sequences involved in gene regulation in many biological processes and diseases. The lack of a complete comprehension of their biological functionality, especially in a genome-wide scenario, has demanded new computational approaches to annotate their roles. It is widely known that secondary structure is determinant to know RNA function and machine learning based approaches have been successfully proven to predict RNA function from secondary structure information.

Here we show that RNA function can be predicted with good accuracy from raw sequence information without the necessity of computing secondary structure features which is computationally expensive. This finding appears to go against the dogma of secondary structure being a key determinant of function in RNA. Compared to recent secondary structure based methods, the proposed solution is more robust to sequence boundary noise and reduces drastically the computational cost allowing for large data volume annotations.

Scripts and datasets to reproduce the results of experiments proposed in this study are available at: https://github.com/bioinformatics-sannio/ncrna-deep

## Introduction

Recent advances in whole transcriptome sequencing have led to the discovery of novel functional non-coding transcript elements classified into miRNA, siRNA, piRNA and lncRNAs. In the past considered as dark matter, they are recognized nowadays to play key roles in gene expression regulation in many biological processes and diseases [1]. The functional characterization of ncRNAs at wide scale is currently one of the main challenges of modern genome biology as – compared to protein coding RNAs – they are usually less conserved and expressed.

The consolidated evidence that the function of protein coding sequences is strongly associated with the folded secondary and tertiary molecular structure leads to suppose that the secondary structure is a key factor to determine the function of non-coding RNA sequences [2]. Recently, several machine learning based approaches have been successfully proven to predict RNA function (Rfam family) from secondary structure information.

Comparative sequence-based approaches, such as BLAST, are computationally very efficient, but exhibit high false negative rates, as they are not able to detect conserved secondary structures. Folding approaches, such as GraPPLE [3], ignore nucleotide composition, are computationally expensive, and incur in high false positive rates, as sequence information is not taken into account. Approaches that combine both structural and sequential information are preferable for a better tradeoff between false positives and false negatives. To this aim, INFERNAL adopts a stochastic context-free grammar to capture position-specific conservation and incorporates the RNA secondary structure information directly into the mode [4]. A significant improvement with respect to INFERNAL has been obtained with EDeN, a machine learning method that adopts a graph kernel to model the RNA secondary structure input representation [5]. Comparable results have been obtained with nRC, a deep learning approach based on features extracted from secondary structure [6], and RNAGCN, based on a graph convolutional network built on RNA folding data [7].

In this paper we show that non coding RNA function can be predicted with good accuracy just from raw sequence information without the necessity of computing secondary structure features which is known to be computationally demanding. Besides the advantage in terms of computational time this finding poses a question against the dogma of secondary structure being a key determinant of function in RNA. Evidence shows that with a 3 layer Convolutional Neural Network (CNN) the sequence alone is enough to predict the function of an RNA. Moreover, compared to recent secondary structure based methods, the proposed solution is more robust to sequence boundary noise and is able to reject effectively non-functional sequences. The last two advantages together with fast classification speed are essential for large genome annotation.

CNN has emerged as an approach to extract local feature patterns of high-level abstraction from different and sparsely preprocessed data [8, 9]. Then it is likely that high level functional RNA features are directly learned from sequence by a CNN architecture. How such features are related, if they are, to secondary structure features remains an open question.

## Materials and methods

### Datasets

We compare our deep learning based approach against EDeN, nRC, and RNAGCN, the current state-of-the-art. We do not include INFERNAL as its computational cost is prohibitively expensive and, in literature, it has been shown outperformed by EDeN [5]. We design the evaluation experiments considering two datasets: i) a novel dataset composed of sequences extracted from the Rfam database, a collection of non-coding RNA sequences manually grouped in families/classes if they share the same function and have a clear common ancestor [10]; and ii) a public available dataset of ncRNA sequences distributed among 13 functional macro-classes adopted to evaluate RNAGCN and nRC [7], as the authors of RNAGCN do not provide a public available tool.

To build the novel dataset, we started with a set of 197922 sequences distributed among 41 classes. Sequences encoded with letters different from canonical A, T, C, or G were excluded to simplify computation. This is not a limitation as they constitute a very small subset from the total (∼5 out of 1000). We removed classes that can be strongly predicted only by sequence length. To detect such classes we performed a 10-fold cross validation of a C5.0 decision tree algorithm trained only with sequence lengths. The algorithm performed overall with an average accuracy of 0.57 (*±*0.001) and Kappa statistic of 0.54 (*±*0.001), while per class performance were strongly variable (average F1 measure ranging between 0.11 and 0.99). We removed two classes, RRF00163 and RF00409, that can be predicted by sequence length with an average F1 measure greater than 0.80. This reduced the number of classes to 39 and the total number of sequences by 13% to 171106. To keep computational time into limit we further removed sequences greater than 150 bases, and, to make each Rfam class sufficiently representative, we excluded classes with less than 400 samples. This conducted to a final set of 162608 sequences distributed among 29 different Rfam classes (Figure 1). Table 1 shows how Rfam classes are distributed among different non-coding macro classes and Figure 2 shows how sequence lengths are distributed among Rfam classes.

**Table 1:** Distribution of downloaded Rfam classes among non-coding macro classes.

| non-coding class | Rfam classes |
| --- | --- |
| snRNA snoRNA | RF00015, RF00016, RF00020, RF00026, RF00066, RF00097, RF00156, RF00560, RF00619 |
| Cis-regulatory | RF00050, RF00059, RF00162, RF00504, RF00557, RF01055, RF01725, RF01739 |
| miRNA | RF00645, RF00875, RF00876, RF00906, RF01059, RF01942 |
| sRNA | RF00019, RF00169, RF01705 |
| Intron | RF00029 |
| rRNA | RF00001 |
| tRNA | RF00005 |

**Fig 1:**
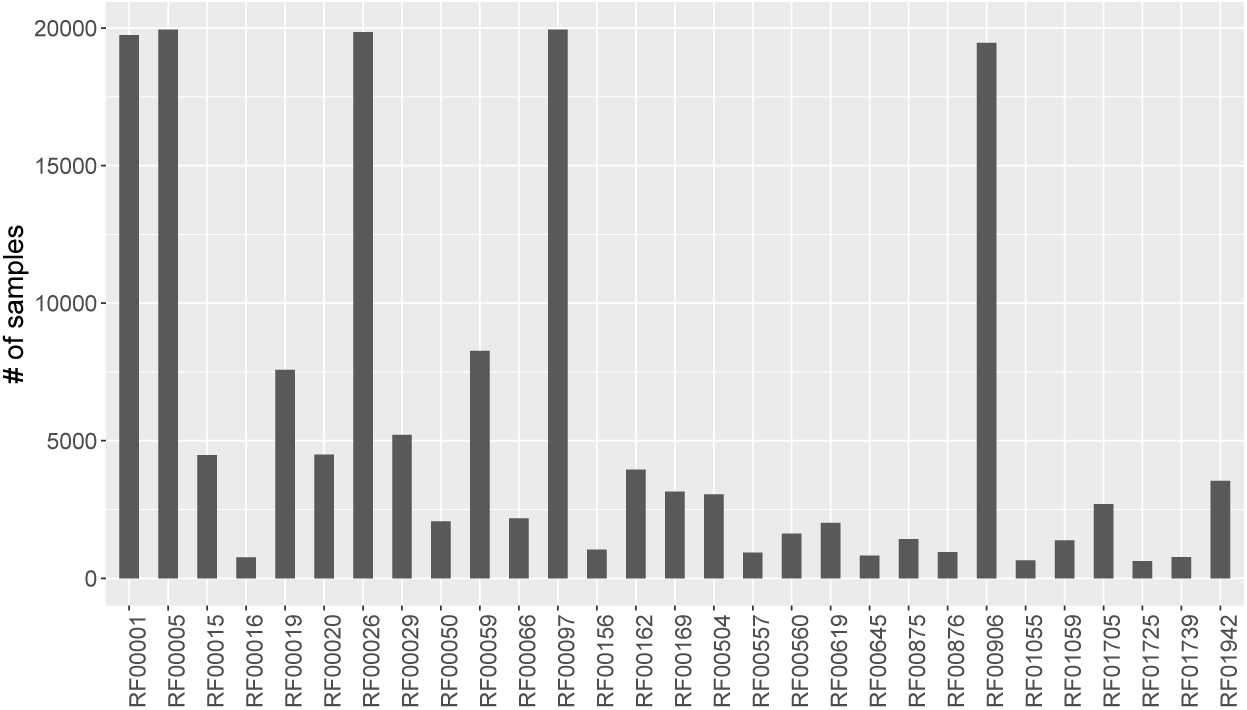
Distribution of sequences among 29 Rfam classes downloaded from Rfam database.

**Fig 2:**
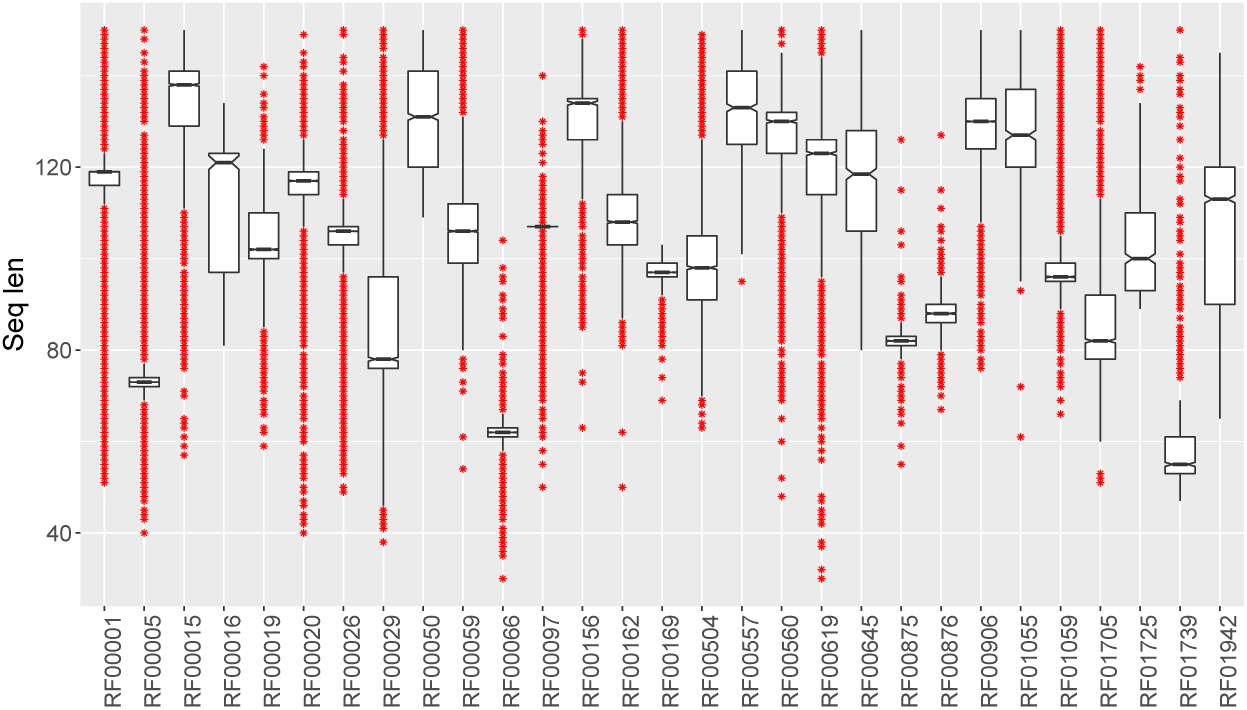
Distribution of sequence lengths among 29 Rfam classes downloaded from Rfam database.

### Input representation of ncRNA sequences

Data representation can strongly affect the performance of machine learning algorithms as they require a good set of hand designed features to work effectively. Instead, the paradigm of deep learning allows in principle to take simple representation of raw data at the lowest (input) layer that is increasingly transformed into abstract feature representations in subsequent layers. However, as deep learning evolved historically around image analysis, the input of a neural network is typically a matrix which has the intrinsic property to completely preserve pixels locality.

In genomics, as the input is a sequence, typical k-mer representation is able to capture the proximal composition of each nucleotide position. This allows to learn local patterns of small nucleotide sequence motifs, such as binding sites, but in principle it may not be suited to detect complex spatial patterns of RNA sequences that fold into a 3-dimensional structure, where also distant nucleotides could interact. So it may be necessary to introduce alternative input representations that allow to map linear structures into bi- or tri-dimensional structures where such patterns could be detected effectively. Current literature methods basically rely on 3-dimensional secondary structure features predicted using popular RNA folding tools, such as ViennaRNA [11] and iPknot [12]. Although such features have been proven to predict RNA function effectively, they require high computational cost (Table 2). Here we investigate whether less computationally expensive sequence encodings are sufficient to predict the RNA function. Specifically we consider k-mer and space-filling curves, a lightweight input representation that preserves, almost well, space locality.

**Table 2:** Computational cost required to build the input representations of a sequence of length *N*.

| Input representation | Computational cost | Adopted in |
| --- | --- | --- |
| Hilbert | $O(N\sqrt{N})$ | Noviello *et al.*, 2020 |
| Morton | $O(N\sqrt{N})$ | Noviello *et al.*, 2020 |
| Snake | $O(N)$ | Noviello *et al.*, 2020 |
| k-mer | $O(N)$ | Noviello *et al.*, 2020 |
| iPknot | $O(N^5)$ | nRC [6] |
| ViennaRNA | $O(N^7)$ | EDeN [5] and<br>RNAGCN [7] |

K-mer encoding is the most common and basic representation of genomic sequence data adopted in deep learning architectures. It consists of associating a binary vector with every consecutive non overlapping *k* bases. The vector is all zeros except for the *i*-th entry associated with the unique *k* word obtained by concatenating *k* letters from the DNA alphabet (Figure 3). So, for example, a 2-mer encoding of a 100 long sequence produces a sequence of 50 binary vectors of 2^4^ = 16 entries. In our experiments we consider *k* varying from 1 to 3.

**Fig 3:**
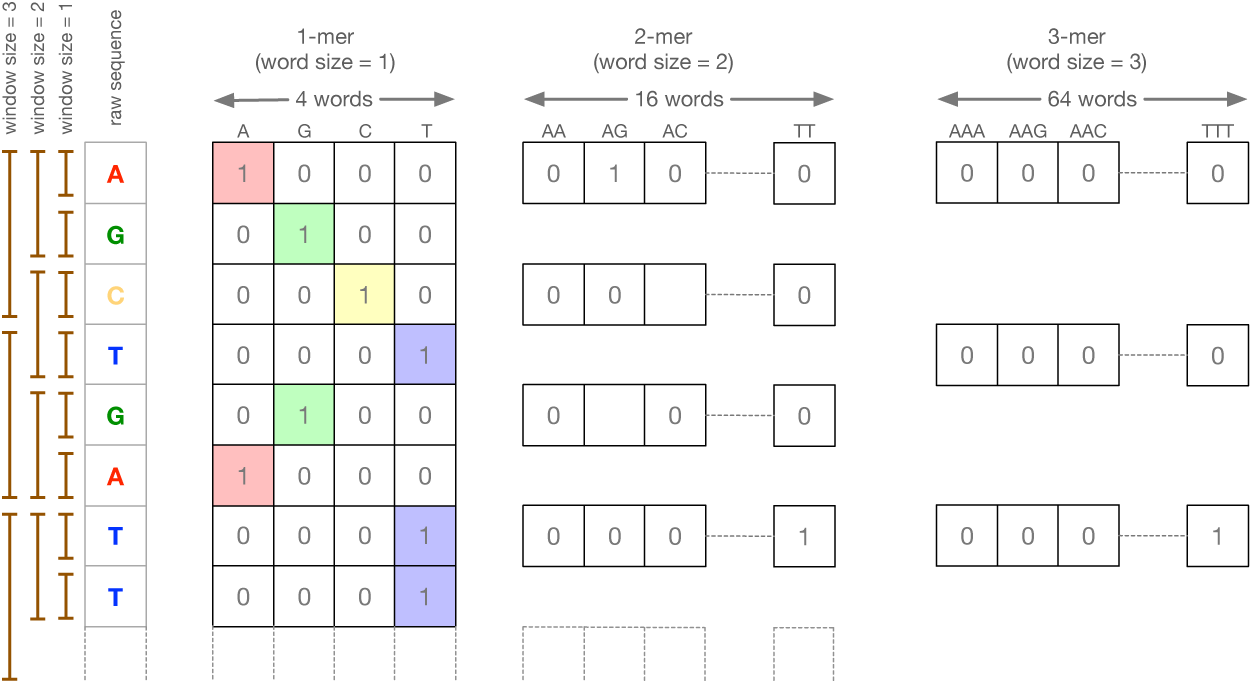
k-mer representation, examples of one, two, and tri-mer encodings.

A space-filling curve is a way to traverse a multi-dimensional space of cell elements where every cell is visited exactly once [13]. Thus, a space-filling curve imposes a linear order of points in the multi-dimensional space that can be mapped to a linear sequence of elements. Different space-filling curves have been proposed, each differing in their way to traversing the multi-dimensional space. We consider three types of 2D space-filling curves: Hilbert [14], Morton [15], and Snake (Figure 4). Each cell is then encoded with a four length binary vector of zeros except for the *i*-th entry associated with the unique DNA letter.

**Fig 4:**
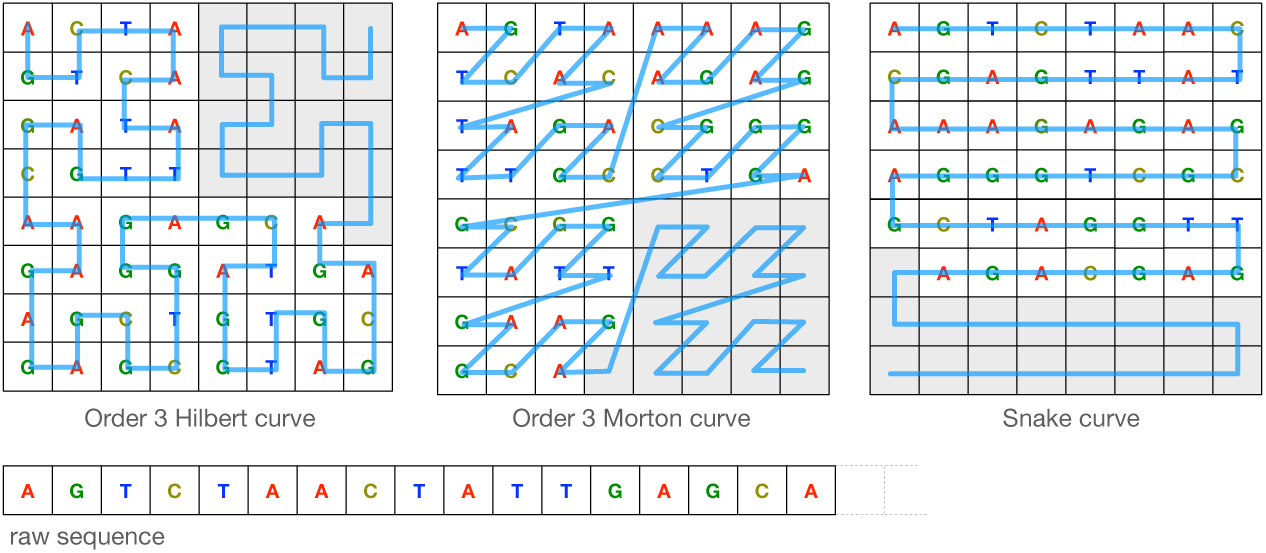
Examples of bi-dimensional space-filling curves. The raw linear 47 base long sequence is encoded into the bi-dimensional space-filling curve depicted in blue. The padding necessary to fill the entire space is depicted in grey.

### Deep network architecture

We adopt the standard deep learning CNN architecture depicted in Figure 5. The network is composed of multiple layers of parametrized kernel convolutions, each composed with: a rectified linear unit (ReLU) activation function to reduce the effect of gradient vanishing, a max-pooling layer to reduce the size of output, and a 50% drop-out layer to reduce overfitting [16]. We consider an increasing number of CNN layers (ranging from 1 to 3), while the dimension of convolution layers is 1D for k-mer input encodings and 2D for space-filling curve encodings. Input sequence representations are first encoded into binary vectors, where each entry corresponds to a CNN channel, and then padded to the maximum dimension allowed for that representation (Table 3). We consider three padding criteria: i) *random*, where vacant cells are filled with random symbols; ii) *constant*, where vacant cells are filled with a constant symbol drawn from the DNA alphabet; and, iii) *new*, where vacant cells are filled with a new symbol not belonging to the DNA alphabet.

**Table 3:** Maximum dimension allowed for each input representation and sequence of at most 150 nucleotides. Dimensions of Hilbert and Morton spaces are the lowest powers of two greater than 150, while the dimension of Snake can be simply obtained consider the ceiling of 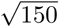.

| Input representation | Maximum dimension allowed |
| --- | --- |
| Hilbert | $16 \times 16$ |
| Morton | $16 \times 16$ |
| Snake | $13 \times 13$ |
| k-mer | $150/k$ |

**Fig 5:**
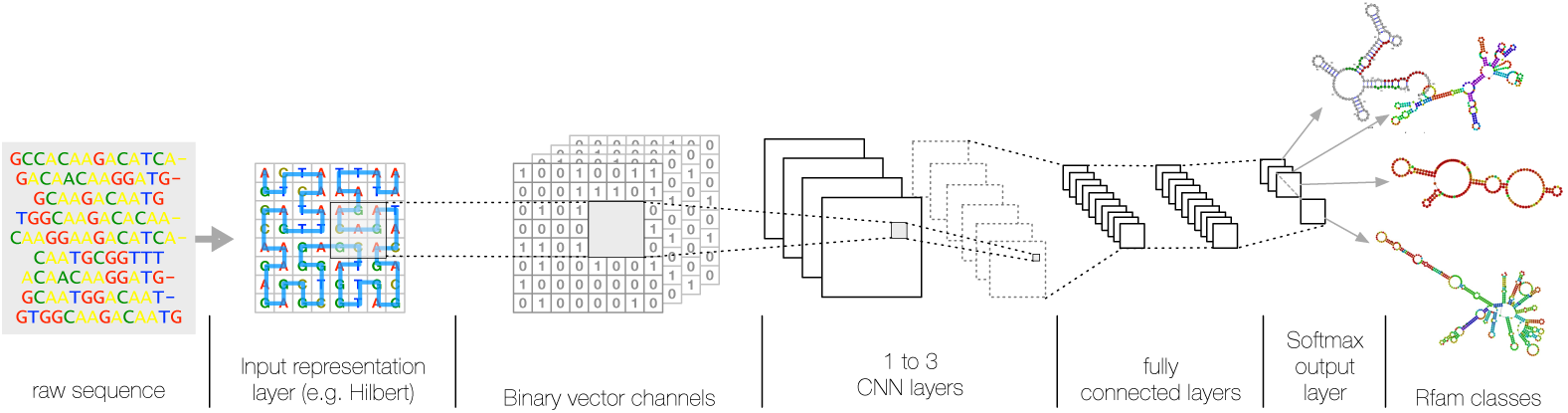
A graphical representation of the deep learning architecture. The raw RNA sequence is first encoded into an input layer representation (e.g. Hilbert filling-curve), then up to 3 convolution layers with rectifier activation followed by max-pooling layers perform the learning steps of sub-sequences with functional properties. Finally, two dense layers of rectified linear units are added to reduce data dimension to a softmax multi-class classification output layer.

We set empirically the kernel size to 3 and the number of filters at each *i*-th layer to 32 · 2^*i*^. The architecture is completed with a flatten layer, to turn spatial features into a vector, two dense layers (respectively of 1000 and 500 nodes), and a softmax output to achieve multi-class classification. For the training step, we adopt Adam [17] as optimization algorithm and categorical cross-entropy loss function, suitable for multi-class classification problem [18].

### Experiment setup

We considered the ncRNA functional annotation task as a multi-class problem where each class is a collection of functionally related ncRNAs. Accuracy and Kappa statistic are adopted to estimate the overall prediction performance, while per class prediction capability is estimated with weighted F1-measure as more informative in highly unbalanced datasets.

To test for the generalization capacity of the algorithm, we split each Rfam class in three random subsets: train (80%), validation (10%), and test (10%). Validation set was used only to tune the hyper-parameters of the learning algorithm, while test set was used to estimate the predictive performance. To limit the bias due to an overpresence of very similar homologous sequences in random splits, we ensured that for each class all sequences in validation and test sets have a similarity - computed in terms of normalized Hamming distance - less than 0.50 with any other sequence in the training set.

Following the experimental assessment conducted in [5] we assess the prediction performance also under the uncertainty of where ncRNA sequence starts and ends. This could happen, for example, with noise coming from next generation sequencing. We added to each sequence a varying *boundary noise*, consisting of a random number of nucleotides at the beginning and the end of a sequence preserving the nucleotide and di-nucleotide frequency of the original sequence [19]. We consider the length of the added noise varying among 0%, 25%, 50%, 75%, 100%, 125%, 150%, 175%, and 200% of the original sequence length.

Moreover we test the rejection capability of the algorithm, *i*.*e*. the behaviour of the algorithm if presented with non-functional RNA sequences – sequences that do not belong to any of the considered classes – or with uncertain sequences. Recently it has been shown that excluding uncertain samples from test set can drastically improve model performance [20, 21]. To this aim we adoped Monte Carlo Dropout to estimate the classification uncertainty of a test sample and decide whether reject or not the sample. We trained the 3 layer CNN architecture with the training set and performed Monte Carlo dropout during test time. Monte Carlo dropout consists to use *N*_*mc*_ different dropout versions of the trained model on the same test sample [20]. In each version, *i* = 1, …, *N*_*mc*_, a random set of nodes is deleted allowing to obtain a discrete probability distribution *p*_*ik*_ among all class values, *k* = 1, …, *C*. From such a distribution the uncertainty of classification can be estimated in different ways [20, 21].

In our experiment we adoped *N*_*mc*_ = 50 and evaluated two uncertainty estimators: *Information Entropy* and *Top Difference*. Information entropy is defined as:

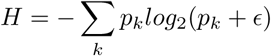

where 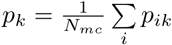, is the mean over all predicted probabilities for a class *k*, and *ϵ* is added for numerical stability. The *Top Difference* is defined as the difference between the two top, in average most probable, predicted classes *k*_1_ and *k*_2_, calculated as:

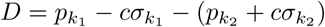

where *σ*_*k*_ is the standard deviation of the discrete probability distribution *p*_*ik*_ among *i*, and *c* a costant we set to *c* = 0.6.

We evaluated the capability to predict functional vs. non-functional RNA sequences plotting the ROC curve of each estimator on a doubled test set obtained by adding to each sequence of the original test set a shuffled version preserving di-nucleotides distribution. Then we evaluated the gain in classification performance on the original test set where uncertain sequences are filtered out considering the following decision thresholds for the estimators: 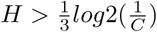 for the Information Entropy estimator and *D <* 0 for Top Difference estimator.

## Results and discussions

### Padding with random symbols affects space filling curve performance

First we evaluated the impact on classification performance of different input sequence representations and padding criteria. To this aim we adopted a 3 CNN layer architecture and evaluated the prediction performance against the novel Rfam dataset. Figure 6 shows the obtained results, in terms of Accuracy (ACC). *k* -mer encodings are not sensitive to padding criteria, while space-filling curve encodings exhibit a significant accuracy drop (∼10-15%) with random padding.

**Fig 6:**
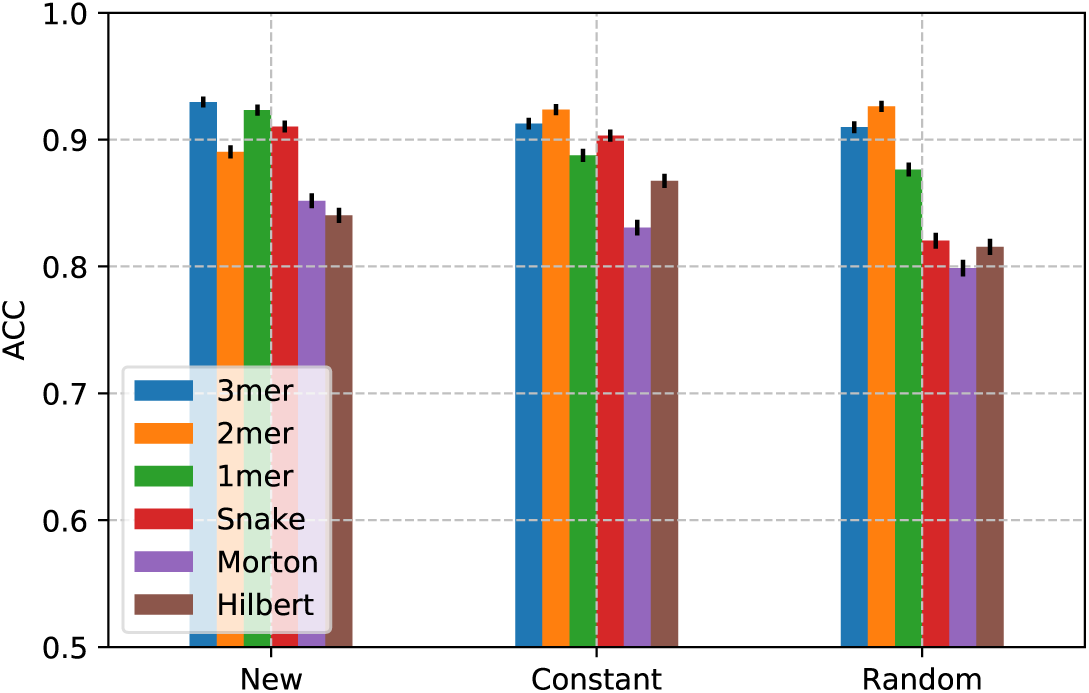
Classification performance in term of Accuracy obtained in the test set with different padding schemas. The deep learning architecture is composed by 3 CNN layers. Confidence intervals are drawn assuming a normal distribution of classification error.

Having in proximity both distal and close elements of a sequence constitute a disadvantage when inputs are filled with random padding, while constant and new symbol padding are less prone to affect overall prediction performance.

### CNN number of layers contributes to performance improvement

Figure 7 shows the impact of neural network depth, codified with the number of CNN layers, on the classification performances. A number of CNN layers equal to zero corresponds to a dense network. According to the results of the previous Section, new symbol has been used as padding criteria and performances were evaluated against the novel Rfam dataset.

**Fig 7:**
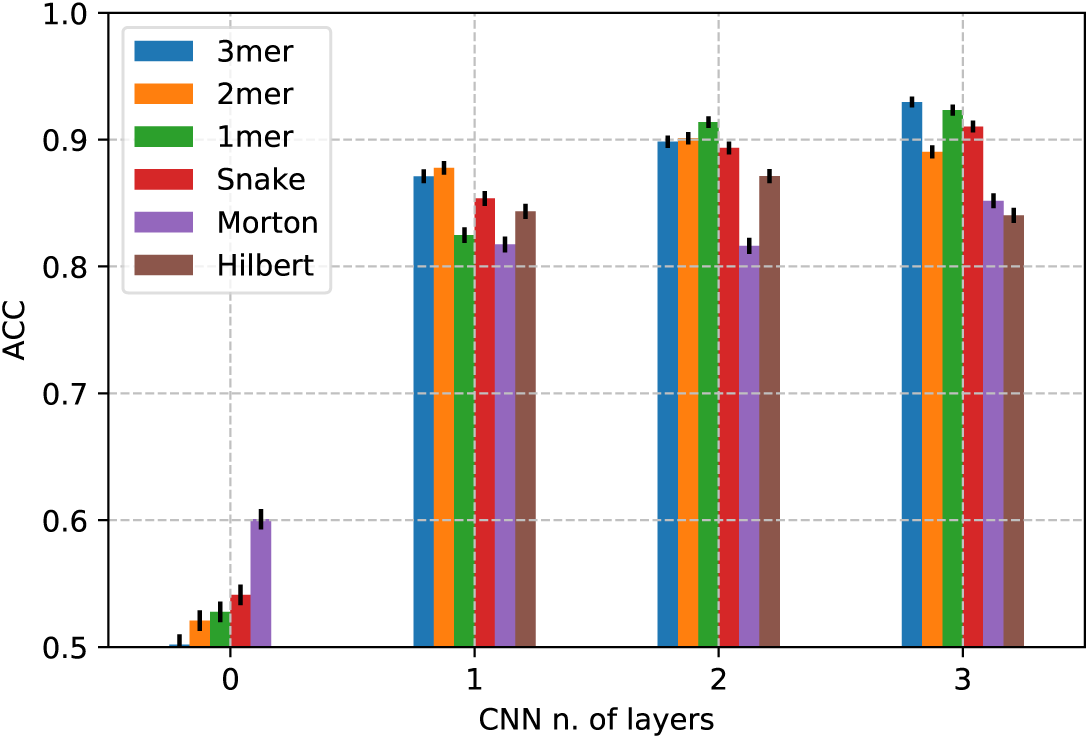
Classification performance in term of Accuracy obtained in the test set with different number of layers using CNNs where inputs are padded with a new symbol. Zero indicates a dense network. Confidence intervals are drawn assuming a normal distribution of classification error.

As expected, the absence of CNN layers strongly affects the learning step resulting in a low rate of accuracy for all the tested input representations. In a dense network, fully connected layers see the data as 1D vectors so it is likely that high level (spatial) relationships and local patterns are not captured. Conversely, increasing architecture’s depth enhances, almost linearly, the learning process of high-level abstract and spatially localized features supposedly connected to RNA function. Adding just one CNN layer increases the prediction accuracy by two fold, advancing it to the range 0.80–0.90 for all input representations. Adding more layers, slightly increases the performance to over 0.90 for some all representations. A significant increment is registered for k-mer and Snake input representations, while adding more layers to Morton and Hilbert representations does not significantly affect the prediction performance. The use of space-filling curves as a proxy for modelling long-range interactions between nucleotides show the worst performance. This does not exactly dismiss the importance of structural effects but poses a question on the necessity to go through the RNA structure to learn RNA functions.

### K-mer encodings are more robust to boundary noise

Figure 8 shows the impact of boundary noise on classification performances against the novel Rfam dataset for each considered input sequence representations. The comparison, in terms of accuracy, with state-of-the-art methods, EdeN and nRC, is also shown. According to the above results, new symbol has been used as padding criteria and three CNN layers as depth of the architecture. At 0% of boundary noise, i.e. original sequences without noise addition, all considered input data representations reach the highest levels of accuracy. With the exception of Morton and Hilbert all the other methods exhibit an accuracy ranging around 0.9. Increasing the percentages of boundary noise a decrement of performance is registered for all methods, more slightly for k-mer representation, while more prominent for spatial-curve representations and even for the state-of-art methods, EdeN and nRC. At 200% of boundary noise the performance of k-mer representations are slightly less than 0.90, in terms of accuracy, while for all others the performance drops to around 0.75.

**Fig 8:**
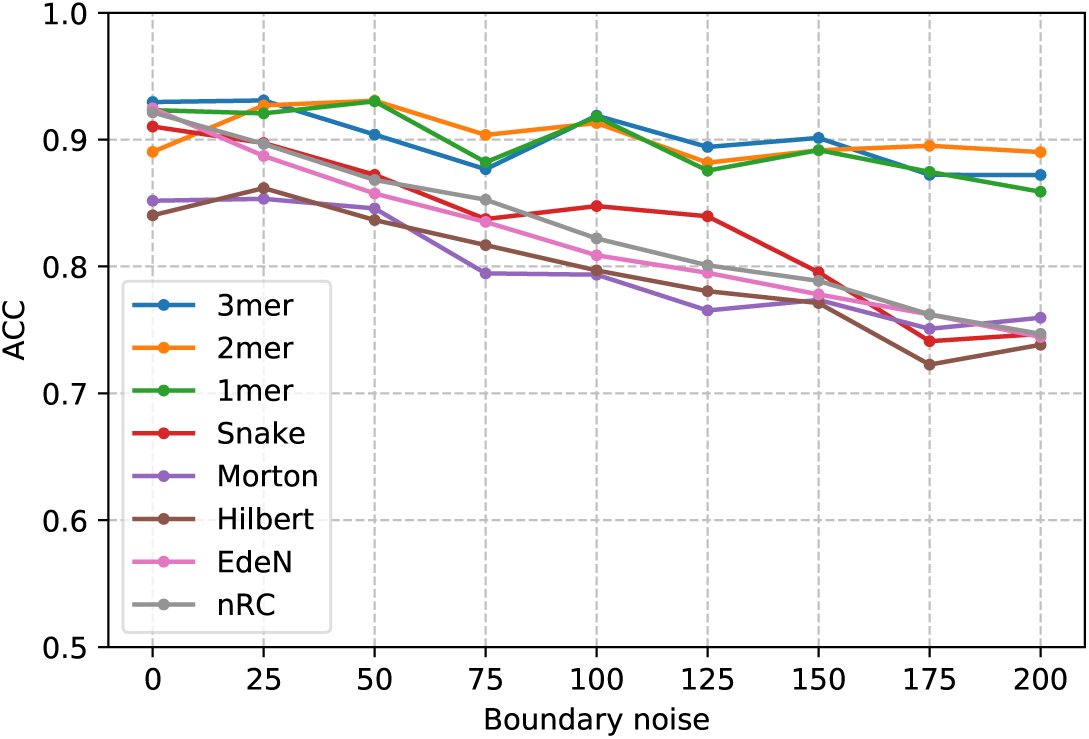
Classification performance in term of Accuracy obtained in the test set at different boundary noise levels. The deep learning architecture is composed by 3 CNN layers and inputs are padded with a new symbol.

Table 4 shows the breakdown of the classification performances of a 3 CNN layers architecture at class level in terms of F1-measure (F1), and its macro and weighted averages. Input sequences are considered with the maximum noise level (200%) and padded with a new symbol. Per class performances with the minimum noise level (0%) are shown in Supplemental Table S1. In terms of weighted averages, the F1-measure of k-mer representations around 7.5% greater than any other method. In terms of macro averages, such increments increase to around 8.3%.

**Table 4:** Per class performance evaluated with a 3 layer CNN network, new padding symbol, and 200% of boundary noise. In each row the maximum value is in boldface.

| Rfam Class | 3mer | 2mer | 1mer | Snake | Morton | Hilbert | EdeN | nRC | Class size |
| --- | --- | --- | --- | --- | --- | --- | --- | --- | --- |
| RF00001 | 0.89 | **0.91** | **0.91** | 0.80 | 0.84 | 0.86 | 0.78 | 0.75 | 3886 |
| RF00005 | 0.87 | **0.88** | 0.84 | 0.81 | 0.76 | 0.73 | 0.77 | 0.82 | 2315 |
| RF00015 | 0.90 | **0.93** | 0.81 | 0.74 | 0.83 | 0.80 | 0.63 | 0.61 | 221 |
| RF00016 | 0.35 | 0.31 | 0.26 | 0.08 | 0.11 | 0.17 | 0.32 | **0.38** | 134 |
| RF00019 | **0.96** | 0.94 | 0.95 | 0.90 | 0.95 | 0.91 | 0.86 | 0.88 | 456 |
| RF00020 | 0.39 | 0.45 | 0.45 | 0.17 | 0.23 | 0.21 | **0.58** | 0.48 | 86 |
| RF00026 | 0.97 | **0.98** | 0.96 | 0.92 | 0.91 | 0.90 | 0.90 | 0.88 | 1813 |
| RF00029 | **0.04** | **0.04** | 0.02 | 0.01 | 0.01 | 0.00 | 0.03 | 0.02 | 5 |
| RF00050 | **0.98** | **0.98** | 0.95 | 0.94 | 0.86 | 0.88 | 0.78 | 0.77 | 378 |
| RF00059 | 0.93 | **0.94** | 0.93 | 0.74 | 0.75 | 0.73 | 0.78 | 0.80 | 1561 |
| RF00066 | 0.69 | 0.58 | **0.84** | 0.64 | 0.63 | 0.67 | 0.76 | 0.76 | 137 |
| RF00097 | 0.96 | **0.97** | 0.96 | 0.94 | 0.93 | 0.94 | 0.72 | 0.81 | 717 |
| RF00156 | **0.71** | 0.65 | 0.62 | 0.32 | 0.37 | 0.39 | 0.41 | 0.40 | 79 |
| RF00162 | 0.69 | 0.73 | **0.82** | 0.46 | 0.45 | 0.28 | 0.58 | 0.48 | 128 |
| RF00169 | 0.89 | **0.91** | 0.83 | 0.63 | 0.80 | 0.75 | 0.66 | 0.59 | 381 |
| RF00504 | 0.89 | **0.92** | 0.89 | 0.53 | 0.61 | 0.71 | 0.79 | 0.80 | 478 |
| RF00557 | **0.88** | 0.87 | 0.86 | 0.59 | 0.66 | 0.64 | 0.48 | 0.35 | 74 |
| RF00560 | 0.84 | **0.86** | 0.71 | 0.36 | 0.78 | 0.57 | 0.54 | 0.45 | 111 |
| RF00619 | 0.43 | **0.54** | 0.40 | 0.27 | 0.44 | 0.37 | 0.30 | 0.24 | 48 |
| RF00645 | **0.82** | 0.77 | 0.85 | 0.23 | 0.27 | 0.21 | 0.51 | 0.66 | 62 |
| RF00875 | 0.93 | **0.95** | 0.91 | 0.85 | 0.42 | 0.57 | 0.46 | 0.66 | 125 |
| RF00876 | **0.45** | 0.36 | 0.40 | 0.55 | 0.34 | 0.27 | 0.27 | 0.39 | 28 |
| RF00906 | 0.78 | 0.76 | **0.81** | 0.80 | 0.76 | 0.71 | 0.76 | 0.78 | 75 |
| RF01055 | **0.79** | 0.77 | 0.76 | 0.67 | 0.58 | 0.57 | 0.47 | 0.46 | 111 |
| RF01059 | 0.69 | **1.00** | **1.00** | 0.90 | **1.00** | 0.23 | 0.56 | 0.25 | 9 |
| RF01705 | 0.84 | **0.86** | **0.86** | 0.72 | 0.71 | 0.66 | 0.81 | 0.81 | 403 |
| RF01725 | 0.26 | 0.46 | **0.51** | 0.22 | 0.30 | 0.25 | 0.48 | 0.36 | 66 |
| RF01739 | 0.25 | **0.43** | 0.18 | 0.26 | 0.17 | 0.12 | 0.30 | 0.37 | 21 |
| RF01942 | **0.95** | 0.91 | 0.91 | 0.91 | 0.79 | 0.82 | 0.85 | 0.86 | 622 |
| Weighed avr | 0.89 | **0.90** | 0.88 | 0.78 | 0.79 | 0.78 | 0.77 | 0.77 | - |
| Macro avr | 0.73 | **0.75** | 0.73 | 0.58 | 0.60 | 0.55 | 0.59 | 0.58 | - |
| Accuracy | 0.87 | **0.89** | 0.86 | 0.75 | 0.76 | 0.74 | 0.74 | 0.75 | - |

A high concordance of per class performance can be observed within the k-mers group, between EDeN and nRC, and within spatial-filled curves. There are classes where all methods are wrong in a similar way, such as: RF00016, RF00020, RF00029, RF00619, RF00876, RF01725, RF01739. Instead, other classes where k-mer, and in some cases spatial-curves, go significantly better than the state-of-art, such as: RF00156, RF00162, RF00169, RF00557, RF00560, RF00645, RF00875, RF01055, RF01059. For all the other classes good performance results are uniformly distributed among all methods. No complementarity among methods can be observed. K-mer representations outperform all other methods, predicting almost correctly 22 out of 29 classes (F1-measure *>* 0.70).

### Monte Carlo Dropout robustly recognizes non-functional RNA sequences and improves prediction performance on non-rejected sequences

Figure 9 shows the performance, estimated in terms of Area under ROC, of rejecting non-functional RNA sequences of two classification uncertain estimators, Information Entropy and Top Distance. Both estimators exhibit similar performance, 0.94 for Information Entropy and 0.92 for Top Distance.

**Fig 9:**
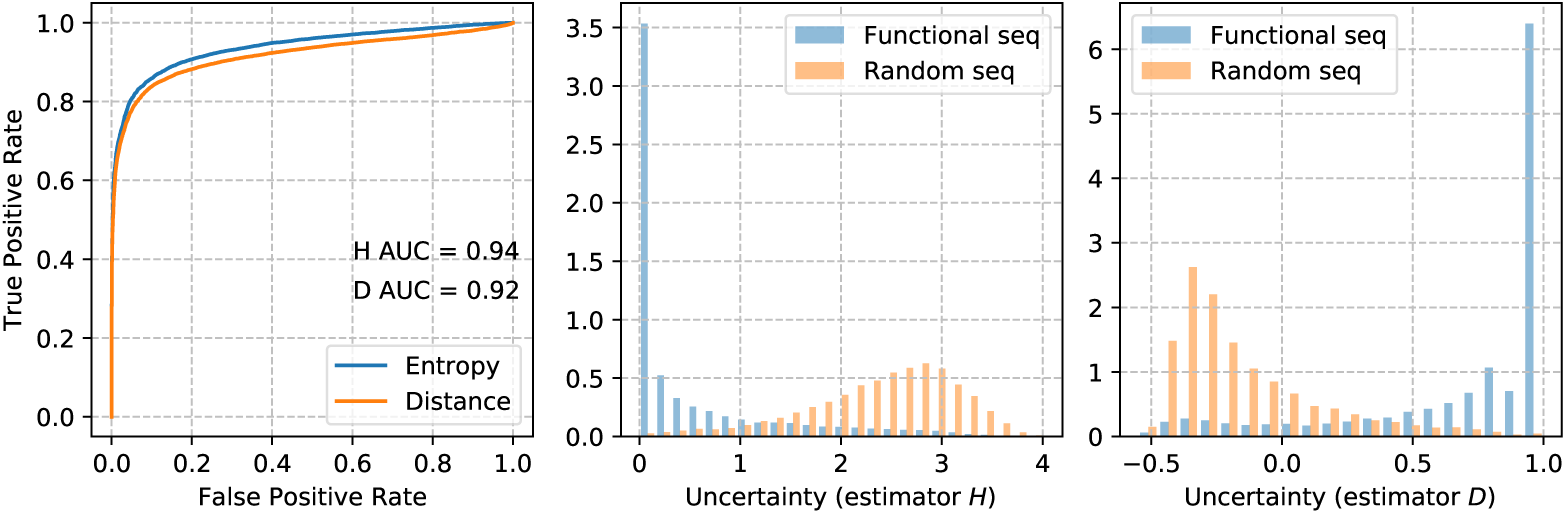
Recognizing non-functional RNA with Monte Carlo Dropout. Sequences are encoded with 1-mer and performance is estimated in terms of Area under ROC (on the left). Figures on the right shows the distributions of functional and non-functional RNA sequences among Information Entropy (H) and Top Distance (D).

Table 5 and Figure 10 show, respectively, the overall and per class performance after Monte Carlo Dropout of uncertain samples encoded with no boundary noise. Overall performance is calculated in terms of Accuracy, Kappa statistic, and Matthew Correlation Coefficient (MCC), while per class performance is calculated in terms of F1-measure. The percentage of dropped samples for each class and the overall percentage of dropped samples are also shown.

**Table 5:** Overall performance improvement, in terms of Accuracy, Kappa, and MCC, after Monte Carlo Dropout of uncertain samples encoded with 1-mer

| Estimator | Approach | Accuracy | Kappa | MCC | % of rejected samples |
| --- | --- | --- | --- | --- | --- |
| Information Entropy | 3mer | 0.97 | 0.97 | 0.97 | 22.05 |
|  | 2mer | 0.99 | 0.98 | 0.98 | 19.81 |
|  | 1mer | 0.99 | 0.99 | 0.99 | 11.16 |
|  | Snake | 0.95 | 0.94 | 0.95 | 26.55 |
|  | Morton | 0.97 | 0.97 | 0.97 | 28.45 |
|  | Hilbert | 0.98 | 0.97 | 0.97 | 33.53 |
| Top Distance | 3mer | 0.97 | 0.97 | 0.97 | 19.69 |
|  | 2mer | 0.98 | 0.98 | 0.98 | 18.09 |
|  | 1mer | 0.99 | 0.99 | 0.99 | 10.85 |
|  | Snake | 0.94 | 0.93 | 0.94 | 25.62 |
|  | Morton | 0.96 | 0.96 | 0.96 | 22.96 |
|  | Hilbert | 0.97 | 0.97 | 0.97 | 27.45 |

**Fig 10:**
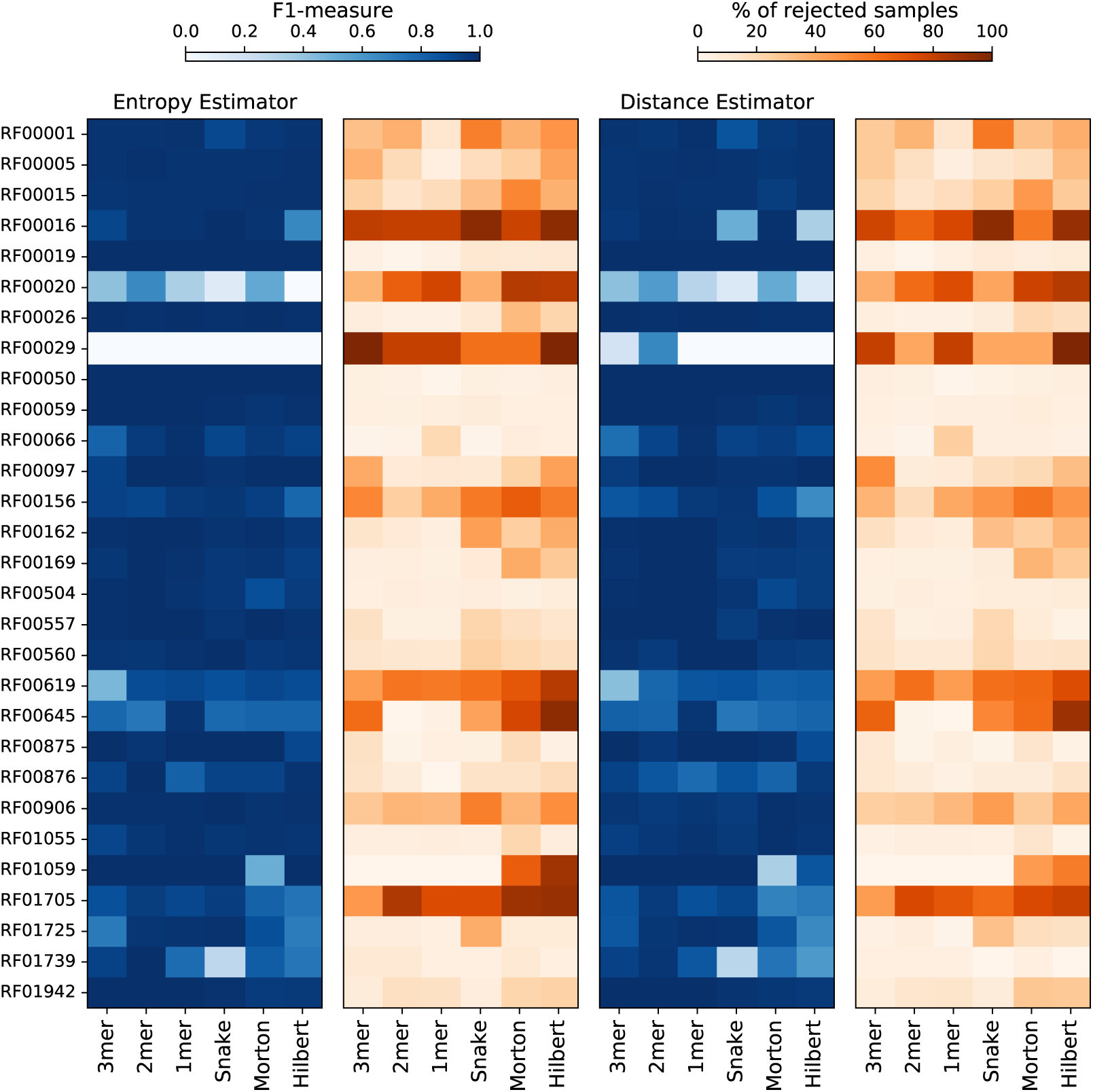
The effect on per class prediction performance of rejecting uncertain samples (1-mer encoded) with Monte Carlo Dropout

For all input representations an overall increment of accuracy can be registered. Comparing results reported in Table 5 with Supplemental Table S1 the following increments can be observed for Information Entropy, 3-mer 4.30%, 2-mer 11.23%, 1-mer 7.60%, Snake 4.39%, Morton 14.11%, Hilbert 16.66%; and the following for Top Distance, 3-mer 4.30%, 2-mer 10.11%, 1-mer 7.60%, Snake 3.29%, Morton 12.94%, Hilbert 15.47%. The highest percentage of dropout samples is registered for Hilbert with Information Entropy (33.53%), while the lowest is registered for 1-mer with Top Distance (10.85%)

The worst per class performance is registered for RF00029 and RF00020 which are also strongly drooped out. Other classes, such as RF00016, RF00619, RF00645, RF01705, gain more performance (see Supplemental Table S1) but are also strongly dropped out.

### Comparison with RNAGCN

The recent proposed RNAGCN method, based on a graph convolutional network, is evaluated against a dataset where ncRNA sequences are classified over 13 functional macro-classes [7]. As the authors of such a method do not provide an executable tool, we were not able to evaluate the proposed method against our novel Rfam dataset containing 30 Rfam classes. So we evaluated our approach against the public available datasets which were originally adopted by the authors of nRC [6]. In particular we choose the dataset called *test13* which is the one that shows best RNAGCN results.

Table 6 reports the results obtained. Surprisingly EDeN exhibits a performance that is significantly lower, in terms of Accuracy, than those obtained against the novel Rfam dataset. Instead with a 3 CNN layers architecture of Figure 5 we obtained performances almost similar to RNAGCN and nRC and almost consistent with those obtained in previous experiments. To test if there is room of improvement we explored alternative CNN architectures. After a thorough empirical evaluation of different architectures we obtained an improved configuration that shows an increment between 5% and 10%, in terms of Accuracy, with respect to the standard architecture adopted in previous experiments (Table 6). The improved architecture is composed of 5 CNN layers interleaved with batch normalization, Leaky ReLU activation, and max-pooling. GaussianNoise to reduce overfitting is added every 2 CNN layers and a dropout rate at 20% is added after the last CNN layer. The network is completed with two dense layers, respectively of 128 and 64 units, to reduce input dimensions, and a final softmax layer for the output class. AMSGrad optimization [22], with a learning rate at 0.0005, has been adopted in the learning step.

**Table 6:**
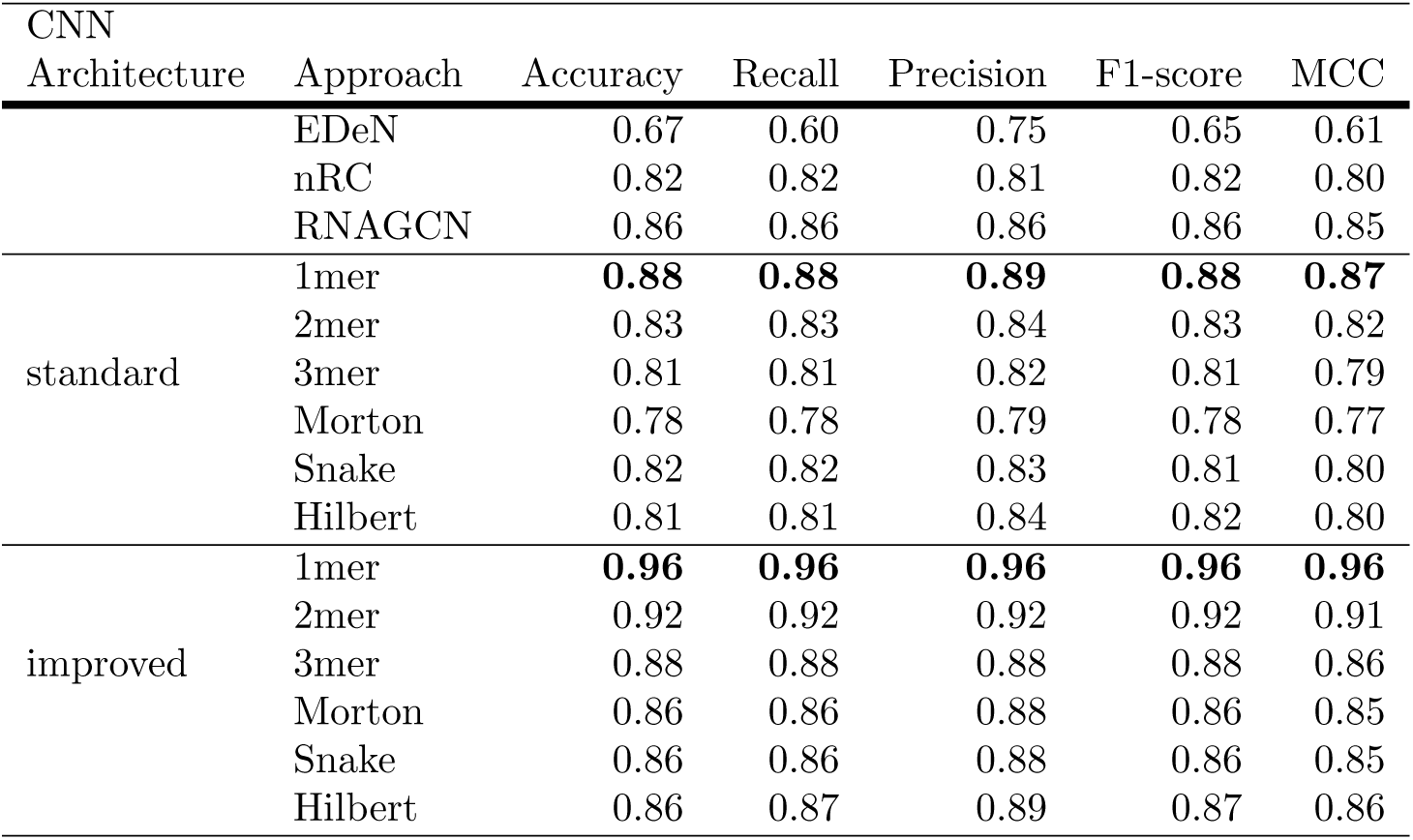
Summary of results on the dataset called *test13* containing 13 non-coding classes. Results for nRC and RNAGCN are taken from [7].

## Conclusion

In this work, we proposed a deep learning approach to classify non coding RNA sequences into Rfam classes. A comparative assessment with the state-of-the-art graph kernel methods shows that the deep learning approach is more robust to boundary noise when k-mer input representations are adopted. CNN number of layers contributes to performance improvement while random padding schema affects negatively space filling curve performance. The deep learning architecture allows for less computational cost input representations than sequence-structure input representations of graph kernel methods. This allows for classification of large scale genomic data and poses and interesting question against the dogma of secondary structure being a key determinant of function in RNA.

The CNN paradigm let us suppose that abstract features associated with RNA functions are effectively learned from simple input representations (i.e. k-mer) and that any further structural encoding in the input representation, such as those carried by space filling curves, does not contribute to performance improvement. To what extend such features are related to secondary structure features remains an open question.

## Acknowledgements

The research leading to these results has received funding from AIRC under IG 2018 ID. 21846 project P.I. Ceccarelli Michele; MIUR PRIN 2017XJ38A4-004; and Regione Campania, POR Campania: Progetto GENOMAeSALUTE (Azione 1.5; CUP: B41C17000080007).

